# Darwinian and Wallacean evolution both explain sexual dichromatism in avian iridescence

**DOI:** 10.64898/2026.11.11.765350

**Authors:** Michaël P.J. Nicolaï, Gerben Debruyn, Rauri C.K. Bowie, Scott V. Edwards, Miguel Carneiro, Liliana D’Alba, Matthew D. Shawkey

**Affiliations:** Biology Department, Evolution and Optics of Nanostructures Group, Ghent University, Ghent 9000, Belgium; Museum of Comparative Zoology, Harvard University, Cambridge, MA; Naturalis Biodiversity Center, Leiden, Leiden, Netherlands; Biology Department, Wybouw lab, Ghent University, Ghent 9000, Belgium; Museum of Vertebrate Zoology and Department of Integrative Biology, University of California, Berkeley, California, USA; CIBIO, Centro de Investigação em Biodiversidade e Recursos Genéticos, InBIO Laboratório Associado, Universidade do Porto, Vairão, Portugal BIOPOLIS Program in Genomics, Biodiversity and Land Planning, CIBIO, Vairão, Portugal

## Abstract

One of the longest-standing arguments in evolutionary biology is that between Darwin and Wallace on the evolution of sexual dichromatism, and which sex changed colour over time. While Darwin argued that males became more colourful over time, Wallace argued that females became less colourful, and numerous studies have lent support to each hypothesis over time. Iridescent colouration provides a particularly interesting system for studying sexual dichromatism because its conspicuousness is often associated with sexual selection and it has evolved multiple times. Here, using a dataset on iridescent colouration of 25,432 patches from 3,151 bird species, we provide one of the most comprehensive tests to date of the evolutionary origins of sexual dichromatism. We show that both hypotheses explain the evolution of iridescent sexual dichromatism in birds, but that Darwin’s hypothesis is more strongly supported with males contributing stronger to evolutionary changes in dichromatism. Furthermore, our analyses support Darwin’s prediction that sexual selection is associated to the evolution of dichromatism, but also Wallace’s prediction that safer nests facilitate more colourful females (and males), and mostly in patches exposed during incubation.

## Introduction

The evolutionary origins of dichromatism have been debated since Darwin (*1*) and Wallace (*2*). At a broad level, their hypotheses differ in which sex is expected to drive the evolution of dichromatism. While Darwin proposed that dichromatism arises primarily through increased male ornamentation (Darwinian evolution), Wallace emphasized female loss of colour (Wallacean evolution). Darwin further argued that elaborate male traits were primarily driven by sexual selection through female preference for conspicuous, and thus colourful, mates (*1*). Wallace argued that sexual selection alone could not account for dichromatism and proposed that natural selection, particularly predation risk during incubation, favoured crypsis in females (2).

Birds provide an ideal system for testing these hypotheses because they are among the most colourful animals, and males are often substantially more colourful than females (*3-7*). In particular, many iridescent species, whose colours change with viewing or lighting angle, exhibit pronounced sexual dichromatism, with males usually displaying more intense colouration than females (*3-7*) (Figure 1A). Within birds, support for each hypothesis has been mixed. Some studies suggest that female plumage colours are more evolutionarily labile and change more frequently than male plumage colours (5). Comparative analyses also support a contribution of female colour evolution to sexual dichromatism in some clades (8-10), but not others (*3, 4*). However, a broad and global comparative assessment across birds, including non-passerines, is lacking.

**Fig. 1.**
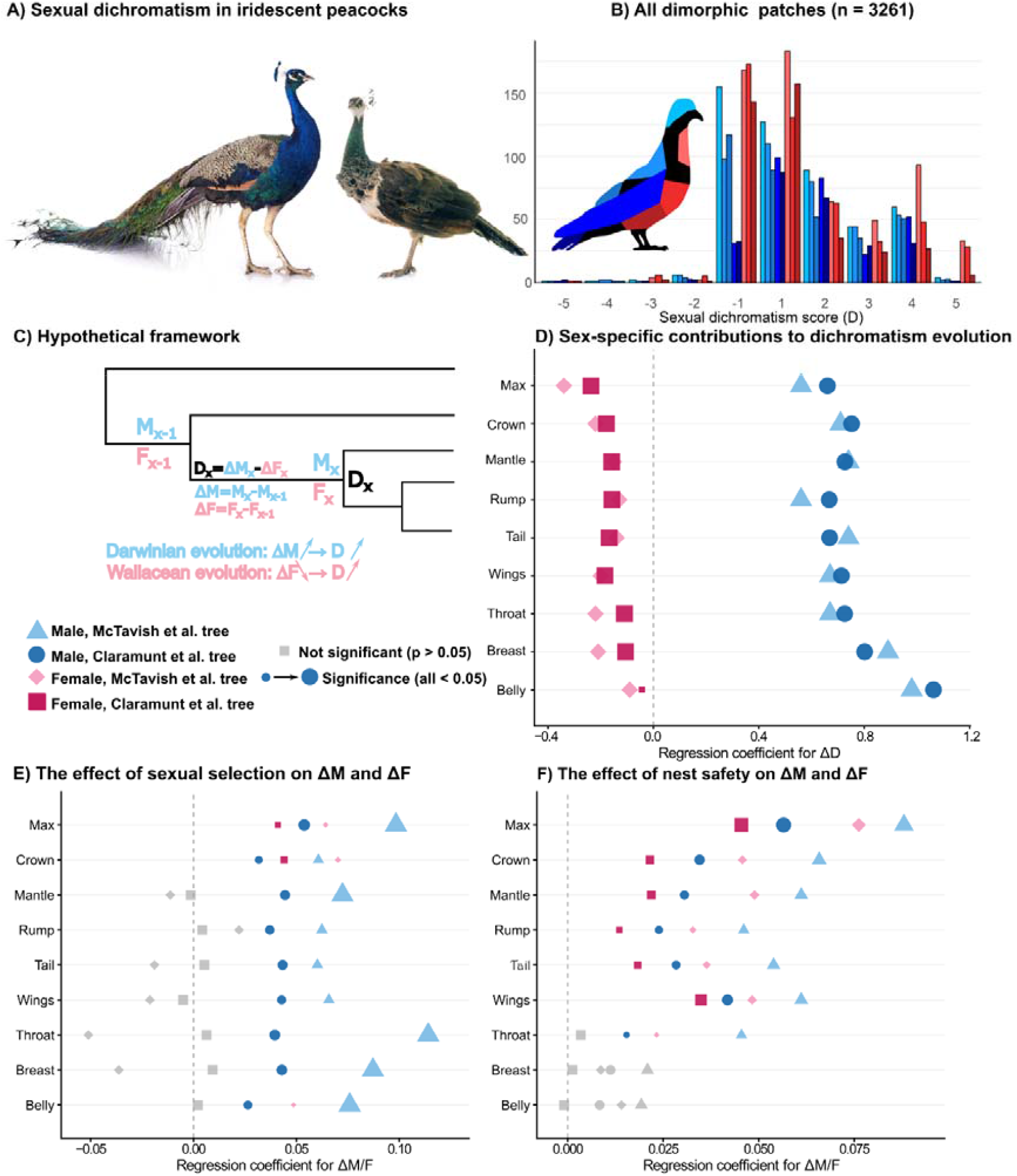
Darwinian and Wallacean contributions to the evolution of sexual dichromatism in avian iridescence. A) Sexual dichromatism in iridescence intensity in Indian peafowl (*Pavo cristatus*). B) Distribution of sexual dichromatism across sexually dimorphic plumage patches (n = 3,261). Positive values indicate greater male iridescence, whereas negative values indicate greater female iridescence. C) Conceptual framework illustrating the two evolutionary routes to increased sexual dichromatism (D): Darwinian evolution through gains in male iridescence (ΔM > 0) and Wallacean evolution through losses in female iridescence (ΔF < 0). D) Associations between evolutionary changes in sexual dichromatism (ΔD) and corresponding changes in male (ΔM) and female (ΔF) iridescence across plumage patches. E, F) Associations between evolutionary changes in male and female iridescence and changes in (E) sexual selection and (F) nest safety. Regression coefficients are shown for two alternative avian phylogenies (McTavish et al. and Claramunt et al.). Blue and pink symbols represent male and female iridescence, respectively. Horizontal position indicates the direction and magnitude of the regression coefficient (β). Zero is shown as a dotted line, and symbol area is proportion to -log10(p).

Here, we use a large, validated dataset comprising 3,151 species, nearly one-third of all extant bird species, for which iridescence intensity data were collected for both sexes. Using these data, we reconstruct the evolution of sexual dichromatism to test the relative contributions of Darwinian and Wallacean processes to the evolution of dichromatism in avian iridescent colouration, and whether the two processes can operate concurrently. We test both scenarios in their broad sense (i.e., males gaining vs females losing iridescence) and in their specific interpretations (i.e. sexual selection favouring colourful males, or natural selection favouring reduced colours in females exposed during incubation). Unlike previous studies, focussing on particular avian clades or more limited phylogenetic sampling, our approach reconstructs the evolutionary history of male and female iridescence separately across Aves, allowing a direct test of the relative contributions of Darwinian and Wallacean processes. In doing so, we provide one of the most comprehensive tests to date of the evolutionary origins of sexual dichromatism in birds.

## Results and discussion

We used a published dataset that scored iridescence intensity from non-iridescent (0) to brilliant iridescence (4) across 25,432 plumage patches from 3,151 bird species (*11*). Most patches showed no sexual dichromatism, but where differences occurred (3261 plumage patches), males were predominantly more iridescent than females (χ^2^ tests, p < 10□□ across regions; Fig. 1B). Dichromatism was greatest in the wings, tail, and throat, consistent with a potential signalling function consisting of patches used in display, or with large visible surfaces (and thus with greater communication, potential)(*3, 12,13*). Strong female-biased iridescence was rare, and in these cases males typically display alternative conspicuous colour mechanisms, such as carotenoid-based colouration (e.g., Many-coloured fruit dove, *Ptilinopus perousii*, with yellow-tailed males and iridescent females).

To distinguish between Darwinian and Wallacean mechanisms, we reconstructed ancestral states of sexual dichromatism (male minus female iridescence intensity) and male and female iridescence independently (*14*), incorporating trait-specific best-fitting evolutionary models and two alternative phylogenies (see Extended Materials and Methods). Increases in dichromatism were more strongly associated with gains in male iridescence (regression coefficients: 0.56 - 1.06) than with losses in female iridescence (-0.34 to -0.04; Fig. 1D). This pattern, that was consistent across plumage patches and both phylogenies, support both Darwinian evolution through gains in male iridescence and Wallacean evolution through losses in female iridescence, although male colour evolution showed consistently stronger associations with dichromatism across plumage patches and phylogenies.

Male associations were weakest for the rump and maximum iridescence score under the McTavish phylogeny (both β = 0.56). The former may reflect reduced signalling opportunities because the rump is often concealed by the wings, whereas the latter could reflect relatively high female iridescence scores, limiting further divergence between sexes. Female associations were strongest for maximum iridescence (β = -0.34) and weakest for the belly (β = 0.002), suggesting that the relative associations of male and female colour evolution with dichromatism vary across plumage patches.

Furthermore, and consistent with Darwin’s predictions, gains in male iridescence were positively correlated with increased sexual selection across all patches, and regardless of phylogeny (Figure 1E). Female iridescence showed little consistent relationship with sexual selection, except potentially for crown and maximum iridescent values across the body, but only when using an alternative phylogeny. These results suggest that although female colour evolution contributes to dichromatism, sexual selection alone cannot fully explain its evolution.

Consistent with Wallace’s prediction that safer nesting environments (defined as a continuous metric combining nest location and structure; Extended Materials and Methods) result in more colourful females, we found that evolutionary increases in nest safety were positively associated with gains in female iridescence across several plumage patches (Fig. 1F). However, similar associations occurred in males, often with larger regression coefficients, suggesting that nest safety is not exclusively associated with female colour evolution. These relationships were strongest in dorsal plumage patches, including the crown, mantle, rump, wings, and tail, whereas ventral regions showed weaker or less consistent associations, which corresponds to patches most likely exposed during incubation. This finding partly agrees with studies linking nesting habits to male colouration (*10*), but contrasts with evidence emphasizing female colouration (*6, 9*).

Differences among studies may reflect variation in taxonomic sampling, colour mechanisms, or analytical approaches (i.e. using only tip-data, see *14*). The different associations of male and female colour evolution are also consistent with a model in which male and female colouration were initially genetically correlated, such that females evolved bright colouration as a correlated response to selection on males before subsequently losing it through natural selection (*15*). Variation in male participation in incubation may also contribute to these patterns, although this remains untested.

Although we find support for both Darwinian and Wallacean evolution, the stronger association between male iridescence gains and dichromatism suggests an asymmetry in their relative contributions. We speculate that this might reflect the unique optical properties of iridescence which can appear both conspicuous and cryptic depending on viewing geometry and lighting conditions, potentially reducing predation pressure without substantially compromising signalling efficacy (*11*).

Thus, while though Darwin’s predictions are more strongly supported, our results suggest that the views of Darwin and Wallace are not mutually exclusive, but instead describe complementary components of the evolutionary dynamics underlying sexual dichromatism; Sexual selection is more consistently associated with male ornamentation, whereas nest safety is associated with colour evolution in both sexes. Whether the stronger contribution of male gains extends to other colour mechanisms and animal lineages remains to be tested.

## Extended Material and Methods

### Quantifying iridescence

We scored the presence of iridescence in 71,536 body patches, from 8942 representative skins of 5755 species of birds (more than 50% of all species) spanning all but 36 genera (98%) from all families, but only retained species for which we had both sexes. For each species we scored the presence and intensity of iridescence in eight patches corresponding to multiple feather regions: crown, mantle, rump, tail, wings, throat, breast and belly. Non-iridescent colours were scored as either melanin-based colours (grey, brown, and black colours), or as non-melanin-based colours. Iridescence was scored by intensity, using four categories adapted from Auber (1957) (*16*) and Durrer (1977) *(17*). Weak iridescence was not readily visible under natural conditions (or in photographs) and generally required close inspection of a specimen (“only close inspection reveals iridescence”; *17*). Moderate iridescence was readily visible under natural conditions (and in photographs), but the plumage retained a predominantly black or dark appearance, with coloured hues becoming apparent only over some feathers or viewing angles (e.g., Rook *Corvus frugilegus*). Strong iridescence was readily apparent under natural conditions (and in photographs), with conspicuous iridescent colour over much of the relevant plumage, although some feathers still appeared black or dark (e.g., the head of Mallard *Anas platyrhynchos*). Brilliant iridescence was conspicuous under natural conditions (and in photographs) across most of the relevant plumage, with most feathers expressing vivid iridescent colour under typical viewing conditions and relatively few appearing black or dark (e.g., many Trochilidae). When in doubt, we used the most conservative score. Importantly, glossy black plumage showing angle-dependent changes in brightness but no discernible change in colour was classified as non-iridescent. These categories were validated using reflectance measurements on a subset of the data (*11*). However, ancestral state reconstructions showed that evolutionary transitions were gradual and rarely skipped intermediate categories (see *11*), consistent with the scale coarsely discretising an underlying continuous trait. Sexual dichromatism was calculated as the difference between male and female scores. Although human visual biases may obscure some chromatic variation (*18*), our categorical approach is likely conservative rather than misleading, as it relies on broad colour classes rather than continuous measures of hue or brightness. Birds can perceive UV signals, and some sex differences may therefore remain undetectable to humans (*18*). To assess this possibility, we examined 229,290 publicly available (*19*) reflectance spectra and found UV peaks in only ∼1% of cases, a small subset of which involved iridescence. Because iridescence, by definition, changes colour with viewing angle and operates over a narrow wavelength range, species exhibiting exclusively UV iridescence are unlikely to have been overlooked. Finally, our classification of sexual dichromatism is based on differences among broad and visually distinct categories, reducing the likelihood of substantial sex misclassification.

### Testing for Darwinian or Wallacean evolution

To distinguish between Darwinian and Wallacean routes to sexual dichromatism, we reconstructed sexual dichromatism (male minus female iridescence score) and male and female iridescence separately. All analyses were repeated using two alternative avian phylogenies, one published by McTavish et al. (*20*) and a second published by Claramunt et al. (*21*).

For each trait, we compared Brownian motion (BM), Ornstein-Uhlenbeck (OU), and Early Burst (EB) models using phylolm (*22*) and selected the model with the lowest AIC. Ancestral states were reconstructed under the selected model using fastAnc (*23*) for BM, GLS-based OU reconstruction for OU, and fastAnc on an EB-rescaled tree for EB using rescale (*24*).

Model comparisons favoured OU for sexual dichromatism across all regions and both phylogenies, except for breast dichromatism under the McTavish phylogeny, for which BM was preferred. In contrast, the best-fitting models for male and female iridescence varied among traits and phylogenies. Under McTavish, BM and OU were favoured for male iridescence, whereas female iridescence additionally showed support for EB. Under Claramunt, OU was favoured for nearly all traits, except male tail iridescence, for which BM was preferred. Complete AIC weights are provided in the accompanying dataset.

For each internal branch, we calculated evolutionary change as the difference between descendant and ancestral node estimates (Δ = descendant − ancestor; Fig. 1C). For each feather patch, we fitted separate linear models relating branchwise changes in sexual dichromatism (ΔD) to corresponding changes in male (ΔM) and female (ΔF) iridescence. Positive associations between ΔD and ΔM were interpreted as consistent with Darwinian evolution, whereas negative associations between ΔD and ΔF were interpreted as consistent with Wallacean evolution.

To test Darwin’s and Wallace’s more specific ecological predictions, we separately reconstructed ancestral states of sexual selection, nest safety, and male and female iridescence using fastAnc, and calculated evolutionary changes along internal branches as described above. We then used generalized least-squares (GLS) models (*25*) to assess whether branchwise changes in male or female iridescence were associated with changes in sexual selection or nest safety. Positive associations between male iridescence and sexual selection were considered consistent with Darwin’s prediction, whereas positive associations between female iridescence and nest safety were considered consistent with Wallace’s prediction. These analyses were performed for each feather patch and the maximum iridescence score across regions.

Nest safety (cf. Wallacean) was scored from 0 to 4 based on nest type and location following Delhey et al. (*26*) (where higher scores correspond to safer nests), while sexual selection was scored from 0 (monogamy with biparental care) to 5 (polygyny/polyandry with uniparental care.

## Acknowledgments

We thank the EON lab for discussing the results. We also thank Annelore Nackaerts from RMCA, Olivier Pauwels from RBINS, Jeremiah Trimble and Kate Eldridge from MCZ for access to their collections. Finally, this work would have been impossible without the multiple generations of scientists that have collected the birds that were used to quantify iridescence.

## Funding

B.A.E.F. (Belgian American Educational Foundation) (M.P.J.N) FWO (Fonds Wetenschappelijk Onderzoek) (M.P.J.N) UGent BOF mandate (M.P.J.N)

European Research Council under the European Union’s Horizon 2020 research and innovation program; grant agreement No. 101000504 (M.C.)

Portuguese Foundation for Science and Technology (FCT, https://www.fct.pt) research contract CEECINST/00014/2018/CP1512/CT0002 (M.C.).

## Author contributions

Conceptualization: M.P.J.N., R.C.K.B., S.V.E., M.C., L.D.A., M.D.S.

Methodology: M.P.J.N., L.D.A., M.D.S.

Investigation: M.P.J.N. with help from G.D.

Visualization: M.P.J.N.

Supervision: R.C.K.B., S.V.E., M.C., L.D.A., M.D.S.

Writing – original draft: M.P.J.N.

Writing – review & editing: all authors.

## Competing interests

The authors declare no competing interests.

## Data and materials availability

All data to reproduce the results are either present in the manuscript or already publicly available. Code will be made publicly available during and after peer review. Up to that point all code is available upon request.

## Notes

### Competing Interest Statement

The authors have declared no competing interest.

